# Evolution of ribosomal internal transcribed spacers in Deuterostomia

**DOI:** 10.1101/112797

**Authors:** Alexandr Dyomin, Valeria Volodkina, Elena Koshel, Svetlana Galkina, Alsu Saifitdinova, Elena Gaginskaya

## Abstract

Sequences of ribosomal internal transcribed spacers (ITSs) are of great importance to molecular phylogenetics and DNA barcoding, but remain unstudied in some large taxa of Deuterostomia. We have analyzed complete *ITS1* and *ITS2* sequences in 62 species from 16 Deuterostomia classes, with *ITSs* sequences in 24 species from 11 classes initially obtained using unannotated contigs and raw read sequences. A general tendency for both *ITS* length and GC-content increase from interior to superior Deuterostomia taxa, a uniform GC-content in both *ITSs* within the same species, thymine content decrease in sense DNA sequences of both *ITSs* are shown. A possible role of GC-based gene conversion in Deuterostomia *ITSs* evolutionary changes is hypothesized. The first example of non-LTR retrotransposon insertion into *ITS* sequence in Deuterostomia is described in turtle *Geochelone nigra.* The roles of mobile genetic element insertions in the evolution of *ITS* sequences in some Sauropsida taxa are discussed as well.

## 1. Introduction

Repetitive ribosomal RNA genes belong to key elements in Metazoan genome. Yet their structure, functioning and evolution still give rise to endless questions. rDNA repeat units correspond to rDNA transcriptional units coding pre-rRNA molecules separated from each other by intergenic spacers (*IGS*). Each pre-rRNA transcriptional unit consists of *18S, 5.8S* and *28S rRNA* genes and separating internal transcribed spacers, *ITS1* and *ITS2* (Fig. 1) (Singer and Berg, 1991). Gene sequences are highly conservative and have been throughly investigated in various animal taxa. Contrarily, spacer sequences are highly variable, both length- and nucleotide content-wise.

**Fig. 1.**
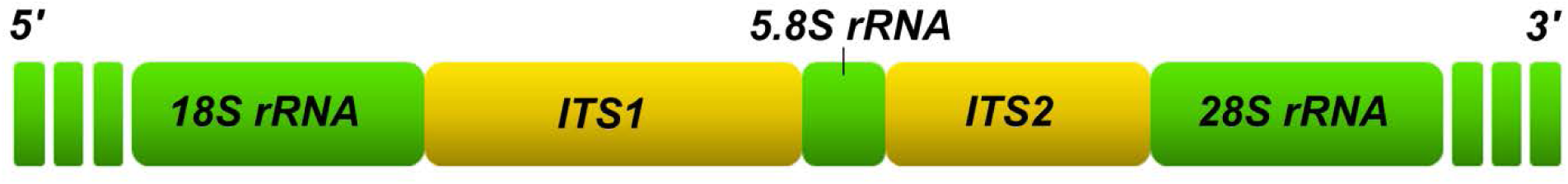
*ITS1* and *ITS2* positions within a rDNA transcription unit

Exploration of *ITS1* and *ITS2* transcribed in pre-rRNA is of special importance. So far their function (presumed regulatory role) so far has remained unclear. However, this high variability resulting from nucleotide substitutions, deletions and insertions turns *ITS*s sequences into a convenient molecular and genetic marker for eucaryotic taxonomy (Yao et al., 2010; Wang et al., 2015). *ITSs* sequences are widely used as phylogenic markers in population and species-grade analysis (Zagoskin et al., 2014). Due to their high variability, *ITSs* sequences can be used in analysis of rDNA repeat unit evolution (Naidoo et al., 2013).

*ITSs* sequences have been throughly analyzed in plants (Chen et al., 2010), fungi (Bellemain et al., 2010) and some Protostomia (Cornman et al., 2015). However, Deuterostoma rRNA structure and evolution, specifically *ITS1* and *ITS2* sequences, are underinvestigated. An exception can be made for some fish and placental mammalian species for which complete sequences of both *ITSs* have been decoded (Coleman, 2013; Wang et al., 2015). *ITS2* structure has been also described for some reptile species (Kupriyanova et al., 2012). For most of other Deuterostomia taxa, *ITSs* sequences are available only for individual species. It is worth noting that NCBI *ITS* annotations for Deuterostomia frequently contain incorrect data on generic identity and size of the spacers (Coleman, 2013; Dyomin et al., 2016). So far no studies *ITSs* structural evolution covering the entire versatility of modern Deutorostomia classes have been conducted.

Such lack of knowledge about Deuterostomia *ITSs* is very likely to be related to their extensive length and high GC-content (Dyomin et al., 2016). These peculiarities of *ITSs* prevent PCR and traditional capillary sequencing application for their primary structural analysis. High GC-content and existence of low complexity regions also impact high-throughput sequencing (HTS) of Deuterostomia *ITSs* sequences. Nonetheless, today HTS findings are the most promising data source for *ITSs* analysis in higher chordate taxa (Coleman, 2013; Dyomin et al., 2016). Due to progress of animal HTS projects (Koepfli et al., 2015), currently data on genomes of over 300 species of various Deuterostomia classes are available (http://www.ncbi.nlm.nih.gov/genome). They are stored in NCBI libraries (Assembly, WGS and SRA) in contig and raw read format. Unannotated contigs and raw reads can contain *ITSs* fragments that can be used for complete sequence assembly (Coleman, 2013; Dyomin et al., 2016).

In this article we have compared *ITS1* and *ITS2* structures of species from 16 Deuterostomia classes. We have identified the common aspects and differences in *ITS1* and *ITS2* of species of various Deuterostomia evolutionary branches. Assumptions on evolutionary patterns of *ITS* sequences in Deuterostomia have been offered.

## 2. Material and methods

Apart from previously published Deuerostomia *ITS1* and *ITS2* sequences, we have compared *ITSs* sequences assembled *de novo* from raw reads and contigs found in NCBI libraries. We have used three data categories: 1) sequences annotated as *ITSs* of individual species in Nucleotide database (http://www.ncbi.nlm.nih.gov/nuccore/); 2) unannotated contig sequences from Assembly database (http://www.ncbi.nlm.nih.gov/assembly/) and 3) unannotated raw read sequences from SRA database (http://www.ncbi.nlm.nih.gov/sra/).

Unannotated sequences containing *ITS*s fragments we identified with the aid of BLAST algorithm (https://blast.ncbi.nlm.nih.gov/Blast.cgi). Fragments of mobile components of *ITSs* sequences were identified in Repbase on on-line basis (http://www.girinst.org/repbase/index.html). *De novo* sequence assembly from contigs and raw reads was performed in Geneious v 9.1.5. (http://www.geneious.com/). Most of *ITSs* Mammalia sequences were borrowed from the assembly published by Coleman (2013). Spacer boundaries were identified with the aid of human rDNA repeat unit annotations (NCBI, Nucleotide: U13369) and chicken rRNA gene cluster (NCBI, Nucleotide: KT445934). Nucleotide structure of the sequences was analyzed in MEGA v6.0 (http://www.megasoftware.net/). The data were processed in Statistica v6.0 (http://www.statsoft.com/).

We have analyzed *ITS1* and *ITS2* performed for 4 types: Chaetognatha, Echinodermata, Hemichordata and Chordata. The outgroup consisted of Ctenophora and Cnidaria species. In Chordata *ITSs* sequences have been analyzed for species of all major classes.

Deuterostomia taxonomy was based on NCBI Taxonomy (http://www.ncbi.nlm.nih.gov/taxonomy/). The analyzed taxa are grouped in the charts based on the phylogenic trees proposed by Dunn et al. (2008), Jarvis et al. (2015) and Hedges et al. (2015).

## 3. Results

### 3.1. ITS sequences search and assembly for analysis

We have analyzed complete *ITS1* and *ITS2* sequences for 62 species (see Table). The Table contains references to the taxonomic status of each of the species, complete GC length and content for both *ITS* sequences and source of *ITS* sequences data for each species. For 32 species, including six outgroup members, we have used NCBI *ITS*s that had been earlier annotated by other authors. These sequences cover the entire analyzed data array on Chaetognatha type and Chondrichthyes and Actinopterygii classes from Chordata. Some sequences earlier annotated in GenBank as *ITS1* and *ITS2* fragments related to the analyzed Deuterostomia species were excluded from the analysis due to low quality of the related assemblies or wrong annotation. The analyzed spacer sequences of 13 mammalian species were earleier described by Coleman (2013). Spacers for 15 species were assembled on the basis of unannotated contigs. For *Petromyzon marinus,* a Cyclostomata species, a combined spacer assembly was performed on the basis of an NCBI annotated sequence (NCBI, Nucleotide: AF061798) and unannotated contig sequences.

**Table.**
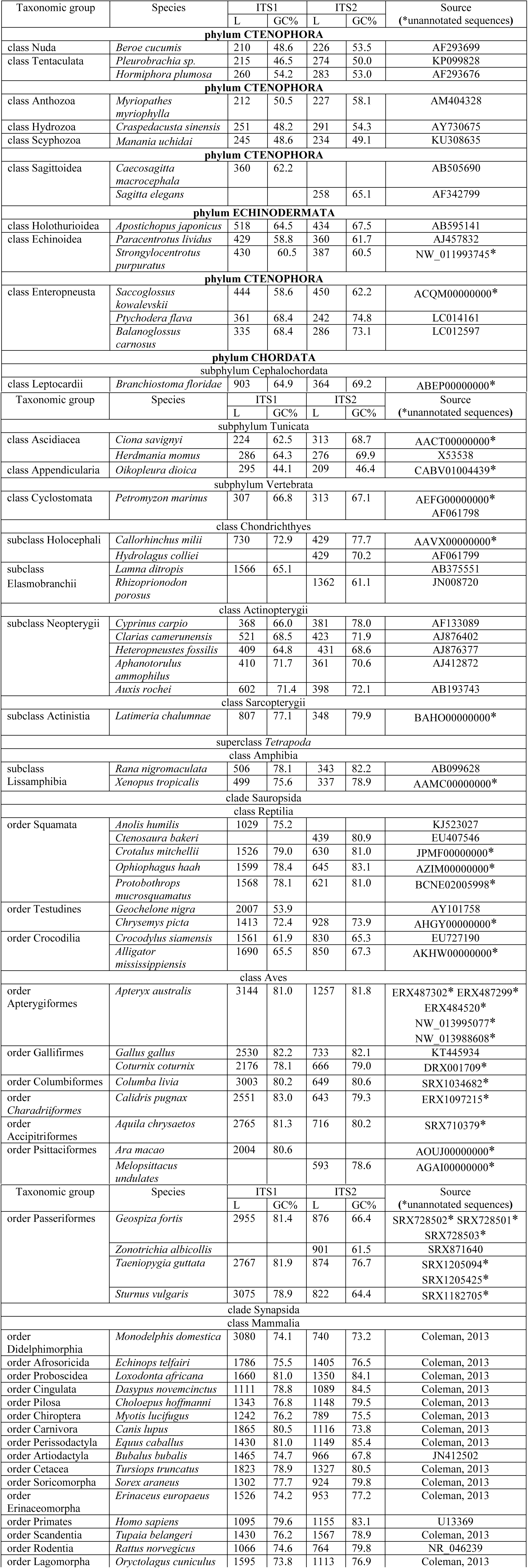
*ITS1* and *ITS2* lengths and GC-content in Deuterostomia representatives

| Taxonomic group | Species | ITS1 |  | ITS2 |  | Source<br>(*unannotated sequences) |
| --- | --- | --- | --- | --- | --- | --- |
|  |  | L | GC% | L | GC% |  |
| phylum CTENOPHORA |  |  |  |  |  |  |
| class Nuda | *Beroe cucumis* | 210 | 48.6 | 226 | 53.5 | AF293699 |
| class Tentaculata | *Pleurobrachia sp.* | 215 | 46.5 | 274 | 50.0 | KP099828 |
|  | *Hormiphora plumosa* | 260 | 54.2 | 283 | 53.0 | AF293676 |
| phylum CTENOPHORA |  |  |  |  |  |  |
| class Anthozoa | *Myriopathes myriophylla* | 212 | 50.5 | 227 | 58.1 | AM404328 |
| class Hydrozoa | *Craspedacusta sinensis* | 251 | 48.2 | 291 | 54.3 | AY730675 |
| class Scyphozoa | *Manania uchidai* | 245 | 48.6 | 234 | 49.1 | KU308635 |
| phylum CTENOPHORA |  |  |  |  |  |  |
| class Sagittoidea | *Caecosagitta macrocephala* | 360 | 62.2 |  |  | AB505690 |
|  | *Sagitta elegans* |  |  | 258 | 65.1 | AF342799 |
| phylum ECHINODERMATA |  |  |  |  |  |  |
| class Holothurioidea | *Apostichopus japonicus* | 518 | 64.5 | 434 | 67.5 | AB595141 |
| class Echinoidea | *Paracentrotus lividus* | 429 | 58.8 | 360 | 61.7 | AJ457832 |
|  | *Strongylocentrotus purpuratus* | 430 | 60.5 | 387 | 60.5 | NW_011993745* |
| phylum CTENOPHORA |  |  |  |  |  |  |
| class Enteropneusta | *Saccoglossus kowalevskii* | 444 | 58.6 | 450 | 62.2 | ACQM00000000* |
|  | *Ptychodera flava* | 361 | 68.4 | 242 | 74.8 | LC014161 |
|  | *Balanoglossus carnosus* | 335 | 68.4 | 286 | 73.1 | LC012597 |
| phylum CHORDATA |  |  |  |  |  |  |
| subphylum Cephalochordata |  |  |  |  |  |  |
| class Leptocardii | *Branchiostoma floridae* | 903 | 64.9 | 364 | 69.2 | ABEP00000000* |
|  |  | L | GC% | L | GC% |  |
| subphylum Tunicata |  |  |  |  |  |  |
| class Ascidiacea | *Ciona savignyi* | 224 | 62.5 | 313 | 68.7 | AACT00000000* |
|  | *Herdmania momus* | 286 | 64.3 | 276 | 69.9 | X53538 |
| class Appendicularia | *Oikopleura dioica* | 295 | 44.1 | 209 | 46.4 | CABV01004439* |
| subphylum Vertebrata |  |  |  |  |  |  |
| class Cyclostomata | *Petromyzon marinus* | 307 | 66.8 | 313 | 67.1 | AEFG00000000*<br>AF061798 |
| class Chondrichthyes |  |  |  |  |  |  |
| subclass Holocephali | *Callorhinchus milii* | 730 | 72.9 | 429 | 77.7 | AAVX00000000* |
|  | *Hydrolagus colliei* |  |  | 429 | 70.2 | AF061799 |
| subclass Elasmobranchii | *Lamna ditropis* | 1566 | 65.1 |  |  | AB375551 |
|  | *Rhizoprionodon porosus* |  |  | 1362 | 61.1 | JN008720 |
| class Actinopterygii |  |  |  |  |  |  |
| subclass Neopterygii | *Cyprinus carpio* | 368 | 66.0 | 381 | 78.0 | AF133089 |
|  | *Clarias camerunensis* | 521 | 68.5 | 423 | 71.9 | AJ876402 |
|  | *Heteropneustes fossilis* | 409 | 64.8 | 431 | 68.6 | AJ876377 |
|  | *Aphanotorulus ammophilus* | 410 | 71.7 | 361 | 70.6 | AJ412872 |
|  | *Auxis rochei* | 602 | 71.4 | 398 | 72.1 | AB193743 |
| class Sarcopterygii |  |  |  |  |  |  |
| subclass Actinistia | *Latimeria chalumnae* | 807 | 77.1 | 348 | 79.9 | BAHO00000000* |
| superclass Tetrapoda |  |  |  |  |  |  |
| class Amphibia |  |  |  |  |  |  |
| subclass Lissamphibia | *Rana nigromaculata* | 506 | 78.1 | 343 | 82.2 | AB099628 |
|  | *Xenopus tropicalis* | 499 | 75.6 | 337 | 78.9 | AAMC00000000* |
| clade Sauropsida |  |  |  |  |  |  |
| class Reptilia |  |  |  |  |  |  |
| order Squamata | *Anolis humilis* | 1029 | 75.2 |  |  | KJ523027 |
|  | *Ctenosaura bakeri* |  |  | 439 | 80.9 | EU407546 |
|  | *Crotalus mitchellii* | 1526 | 79.0 | 630 | 81.0 | JPMF00000000* |
|  | *Ophiophagus haah* | 1599 | 78.4 | 645 | 83.1 | AZIM00000000* |
|  | *Protobothrops mucrosquamatus* | 1568 | 78.1 | 621 | 81.0 | BCNE02005998* |
| order Testudines | *Geochelone nigra* | 2007 | 53.9 |  |  | AY101758 |
|  | *Chrysemys picta* | 1413 | 72.4 | 928 | 73.9 | AHGY00000000* |
| order Crocodilia | *Crocodylus siamensis* | 1561 | 61.9 | 830 | 65.3 | EU727190 |
|  | *Alligator mississippiensis* | 1690 | 65.5 | 850 | 67.3 | AKHW00000000* |
| class Aves |  |  |  |  |  |  |
| order Apterygiformes | *Apteryx australis* | 3144 | 81.0 | 1257 | 81.8 | ERX487302* ERX487299*<br>ERX484520*<br>NW_013995077*<br>NW_013988608* |
| order Gallifirmes | *Gallus gallus* | 2530 | 82.2 | 733 | 82.1 | KT445934 |
|  | *Coturnix coturnix* | 2176 | 78.1 | 666 | 79.0 | DRX001709* |
| order Columbiformes | *Columba livia* | 3003 | 80.2 | 649 | 80.6 | SRX1034682* |
| order Charadriiformes | *Calidris pugnax* | 2551 | 83.0 | 643 | 79.3 | ERX1097215* |
| order Accipitriformes | *Aquila chrysaetos* | 2765 | 81.3 | 716 | 80.2 | SRX710379* |
| order Psittaciformes | *Ara macao* | 2004 | 80.6 |  |  | AOUJ00000000* |
|  | *Melopsittacus undulates* |  |  | 593 | 78.6 | AGAI00000000* |
|  |  | L | GC% | L | GC% |  |
| order Passeriformes | *Geospiza fortis* | 2955 | 81.4 | 876 | 66.4 | SRX728502* SRX728501*<br>SRX728503* |
|  | *Zonotrichia albicollis* |  |  | 901 | 61.5 | SRX871640 |
|  | *Taeniopygia guttata* | 2767 | 81.9 | 874 | 76.7 | SRX1205094*<br>SRX1205425* |
|  | *Sturnus vulgaris* | 3075 | 78.9 | 822 | 64.4 | SRX1182705* |
| clade Synapsida |  |  |  |  |  |  |
| class Mammalia |  |  |  |  |  |  |
| order Didelphimorphia | *Monodelphis domestica* | 3080 | 74.1 | 740 | 73.2 | Coleman, 2013 |
| order Afrosoricida | *Echinops telfairi* | 1786 | 75.5 | 1405 | 76.5 | Coleman, 2013 |
| order Proboscidea | *Loxodonta africana* | 1660 | 81.0 | 1350 | 84.1 | Coleman, 2013 |
| order Cingulata | *Dasypus novemcinctus* | 1111 | 78.8 | 1089 | 84.5 | Coleman, 2013 |
| order Pilosa | *Choloepus hoffmanni* | 1343 | 76.8 | 1148 | 79.5 | Coleman, 2013 |
| order Chiroptera | *Myotis lucifugus* | 1242 | 76.2 | 789 | 75.5 | Coleman, 2013 |
| order Carnivora | *Canis lupus* | 1865 | 80.5 | 1116 | 73.8 | Coleman, 2013 |
| order Perissodactyla | *Equus caballus* | 1430 | 81.0 | 1149 | 85.4 | Coleman, 2013 |
| order Artiodactyla | *Bubalus bubalis* | 1465 | 74.7 | 966 | 67.8 | JN412502 |
| order Cetacea | *Tursiops truncatus* | 1823 | 78.9 | 1327 | 80.5 | Coleman, 2013 |
| order Soricomorpha | *Sorex araneus* | 1302 | 77.7 | 924 | 79.8 | Coleman, 2013 |
| order Erinaceomorpha | *Erinaceus europaeus* | 1526 | 74.2 | 953 | 77.2 | Coleman, 2013 |
| order Primates | *Homo sapiens* | 1095 | 79.6 | 1155 | 83.1 | U13369 |
| order Scandentia | *Tupaia belangeri* | 1430 | 76.2 | 1567 | 78.9 | Coleman, 2013 |
| order Rodentia | *Rattus norvegicus* | 1066 | 74.6 | 764 | 79.8 | NR_046239 |
| order Lagomorpha | *Oryctolagus cuniculus* | 1595 | 73.8 | 1113 | 76.9 | Coleman, 2013 |

Most challenging was *ITSs* analysis in Aves, one of the most progressive classes among Deuterostomia. So far transcribed avian ribosome RNA spacers have remained virtually unknown (Dyomin et al., 2016). An article by Kupriyanova et al., (2012) mentions zebra finch *ITS2* assembly, yet, unfortunately, the complete sequence has not been disclosed. Earlier we annotated in NCBI the first complete assembly of chicken rRNA gene cluster sequence containing both *ITSs* (NCBI, Nucleotide: KT445934). In this survey we have assembled from unannotated contigs one *ITSs* for each of the two Psittaciformes species. We have also been able to decode the complete structure of both *ITSs* for 8 species from 6 avian orders using *de novo* assembly from WGS raw reads. We have also established *ITS2* sequence for *Zonotrichia albicollis* (Passeriformes, Aves).

In total, we have been the first to analyzed *ITS*s structure for 24 species from 11 Deuterostomia classes. All sequences used, including those first assembled in this study, are listed in Supplementary File 1.

### 3.2. ITSs structural trends in Deuterostomia evolution

We have analyzed Deuterostomia *ITSs* structural changes in two evolutionary vectors: from Chaetognatha to Reptilia–Aves (Sauropsida), and from Chaetognatha to Mammalia (Sinapsida). For simplification both these vectors are represented in separate charts designated as Sauropsida line and Synapsida line respectively. In both charts taxa layout is based on their evolutionary proximity. However, closely located taxa are not necessarily ancestors to or descendants of each other.

#### 3.2.1. ITS1 and ITS2 sequence length change

*ITSs* length change in Sauropsida and Synapsida lines is shown in Figures 2 and 3 respectively. By *ITSs* length change nature both lines clearly fall into two parts. Within the left part (Chaetognatha – Amphibia) the average lengths of both *ITSs* are 1.5 times as much as *ITS* lengths in the outgroup: *ITS1* – 419 bp, *ITS2* – 364 bp vs 230 bp and 254 bp in Ctenophora and Cnidaria respectively (Fig. 4). Generally no clear relation between spacer length change and evolutionary development of animal structure can be identified here. The only exception are *ITS* lengths in the analyzed Elasmobranchii (*ITS1* in *Lamna ditropis –* 1566 bp; *ITS2* in *Rhizoprionodon porosus* – 1362 bp), which considerably exceed the average *ITSs* lengths within this group. In terms of *ITS1* length also stand out lancelet (*Branchiostoma floridae*) and coelacanth (*Latimeria chalumnae*) (Fig. 2 and 3). The above exceptions are in line with high variability of *ITS1* (Fig. 4). Yet, generally *ITS1* and *ITS2* average lengths in the first chart part are approximately equal, *ITS1* being only slightly longer than *ITS2* (Fig. 5).

**Fig. 2.**
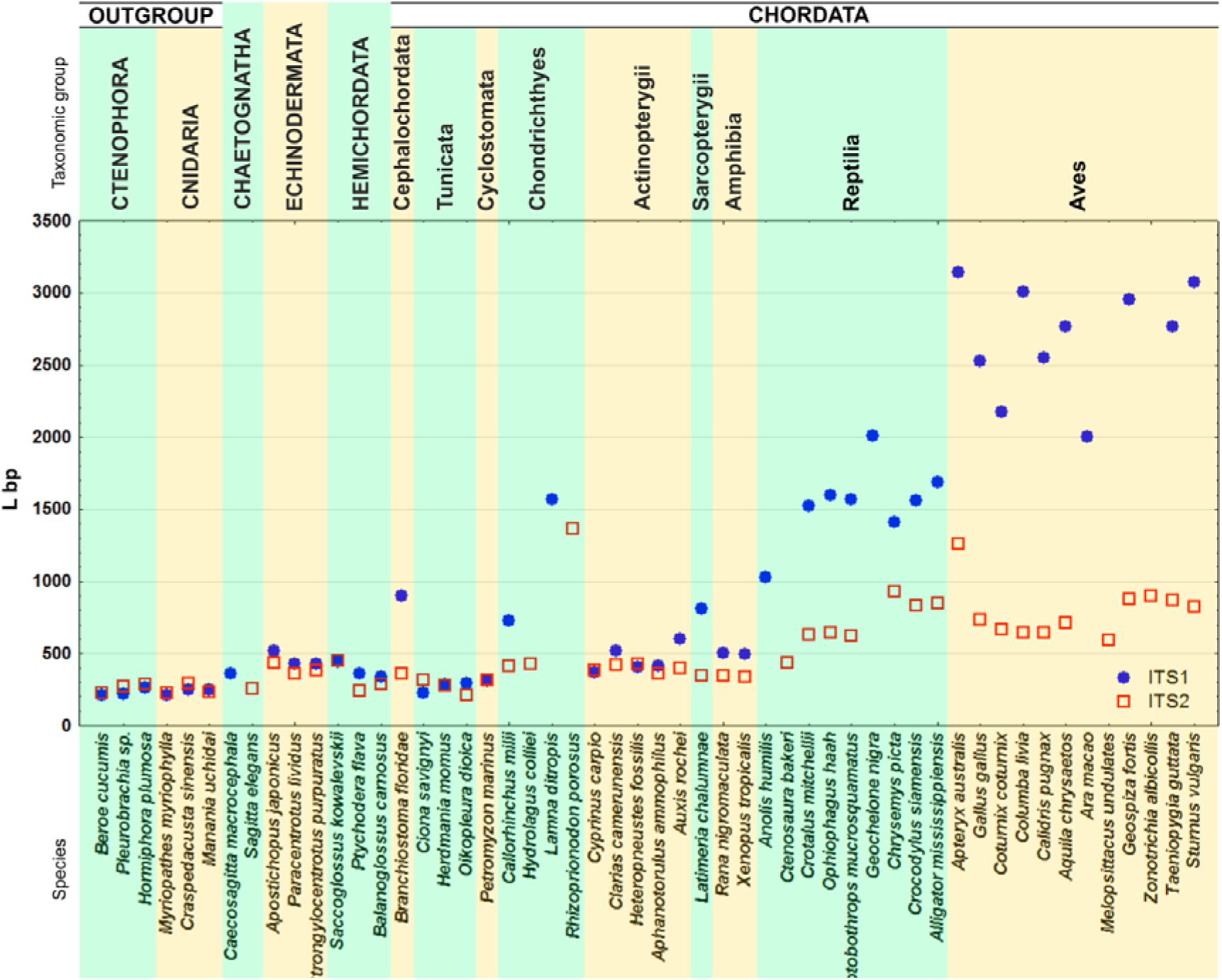
*ITS1* and *ITS2* lengths in Deuterostomia taxa. Sauropsida evolutionary line.

**Fig. 3.**
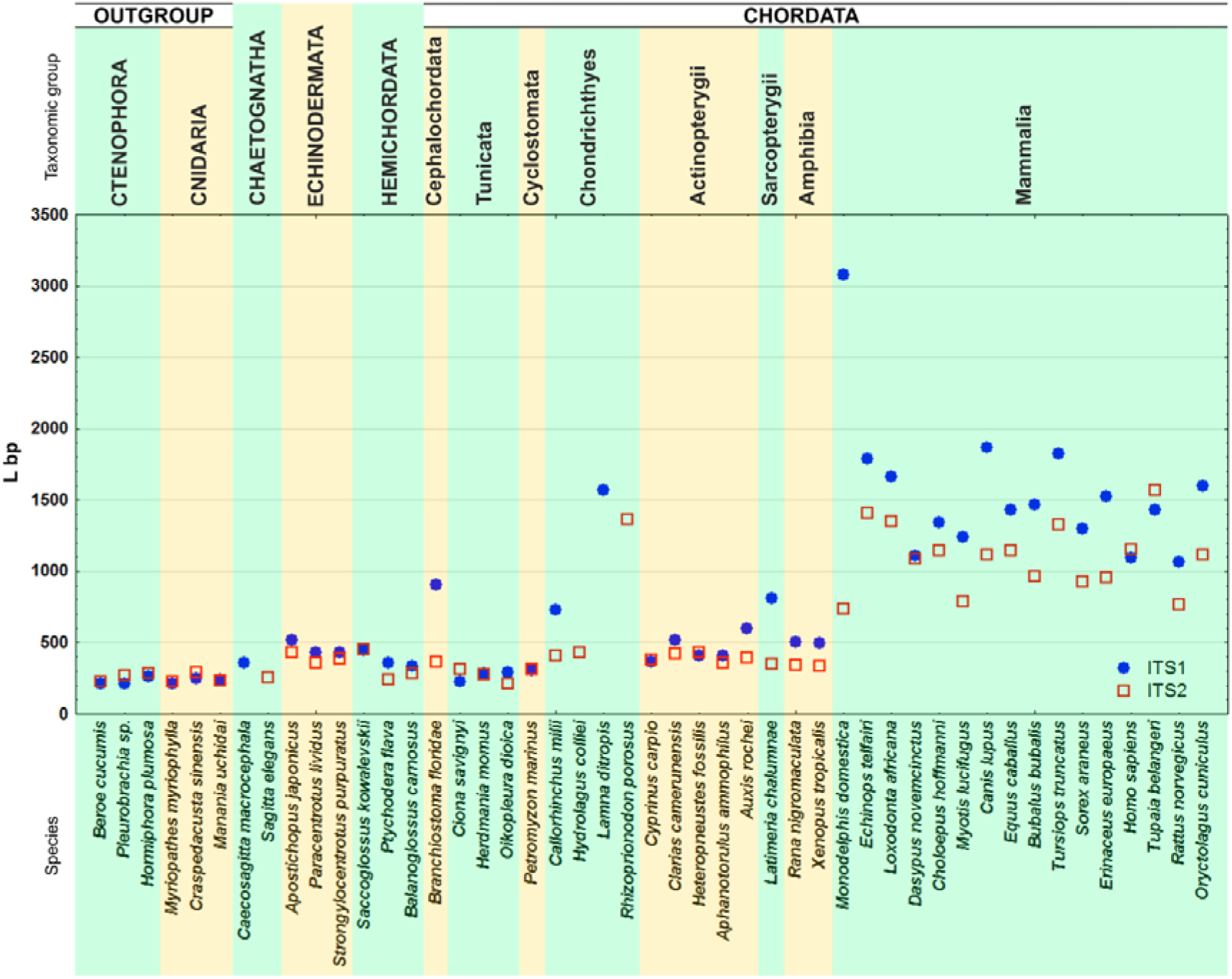
*ITS1* and *ITS2* lengths in Deuterostomia taxa. Synapsida evolutionary line

**Fig. 4.**
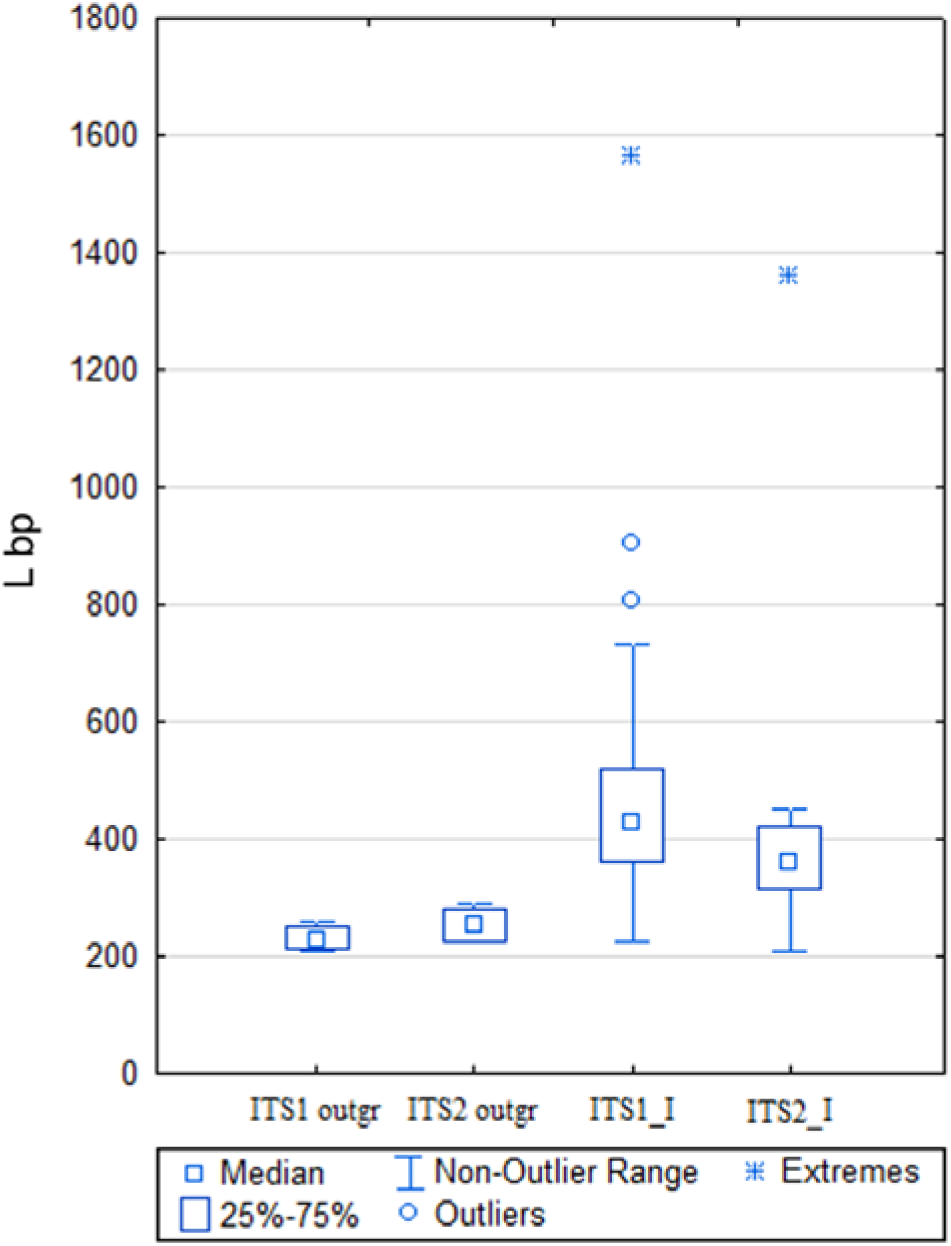
Box plot of *ITS1* and *ITS2* lengths in Chaetognatha-Amphibia. ITS1_I and ITS2_I – *ITS1* and *ITS2* sequence lengths in Chaetognatha–Amphibia species. ITS1_outgr and ITS2_outgr – *ITS1* and *ITS2* sequence lengths in Chaetognatha-Amphibia species

**Fig. 5.**
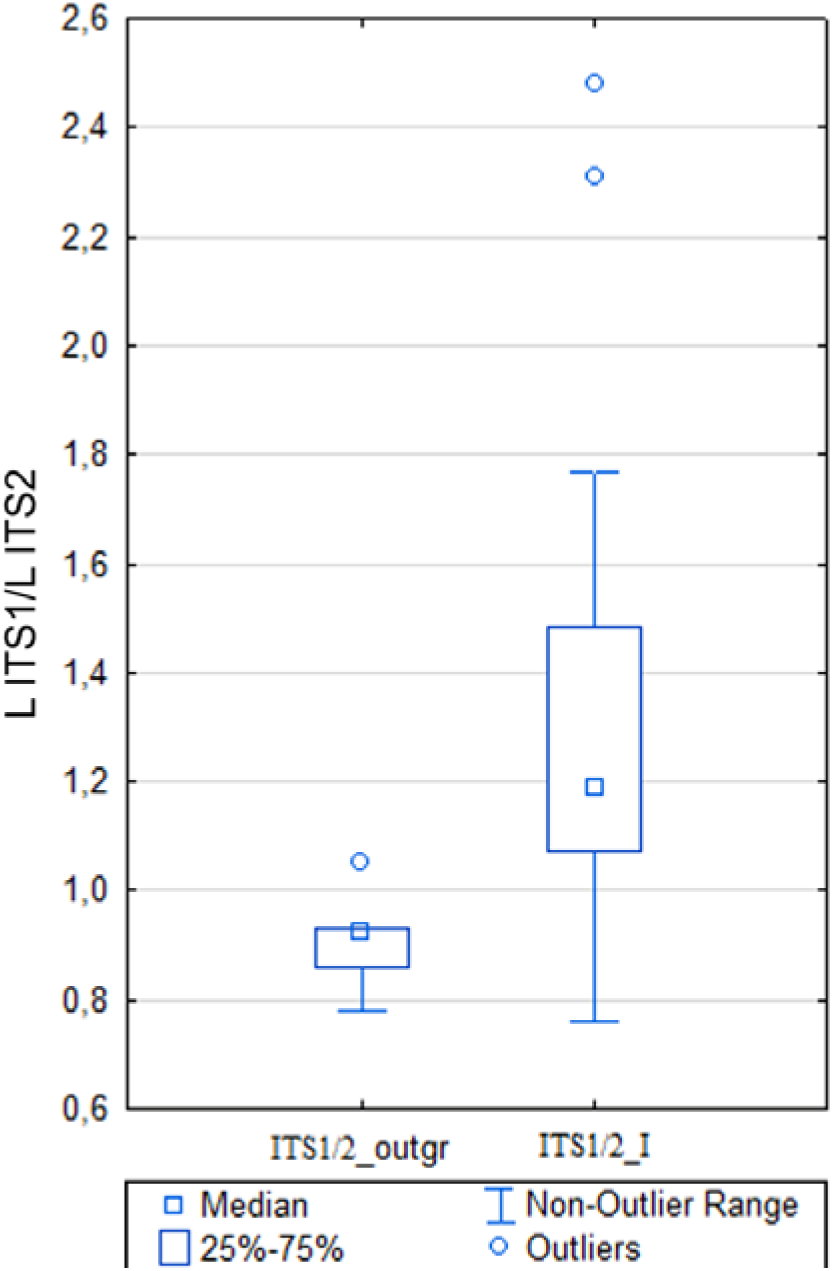
*ITS1* to *ITS2* length ratio in Chaetognatha-Amphibia. ITS1/2_outgr – *ITS1* to *ITS2* length ratio in Ctenophora *u* Cnidaria species; ITS1/2_I – *ITS1* to *ITS2* length ratio in Chaetognatha-Amphibia species.

The right sections of charts 2 and 3 consist of *ITS1* and *ITS2* sequences of Sauropsida (Reptilia, Aves) and Sinapsida (Mammalia) respectively. Both in Sauropsida and Sinapsida both spacers are considerably longer than those of Chaetognata-Amphibia and outgroup. In Sauropsida line *ITS2* length increases smoothly with transition from anamniotes to amniotes (Fig. 2); towards Aves the disproportion in *ITS1* and *ITS2* length increases (Fig. 6). *ITS1* sequence in kiwi (*Apteryx australis*) of 3144 bp is the longest among the analyzed animal species. Taking into account the data reported by Coleman (2013), this value is the ceiling in Deuterostomia. Among Sauropsida, kiwi also features the longest *ITS2* of 1257 bp. *ITS2* length in kiwi, a member of Paleognathae, considerably differs from *ITS2* of the analyzed avian species, sub-class Neognathae, and reptiles whose *ITS2* is 1.5–2 times shorter.

**Fig. 6.**
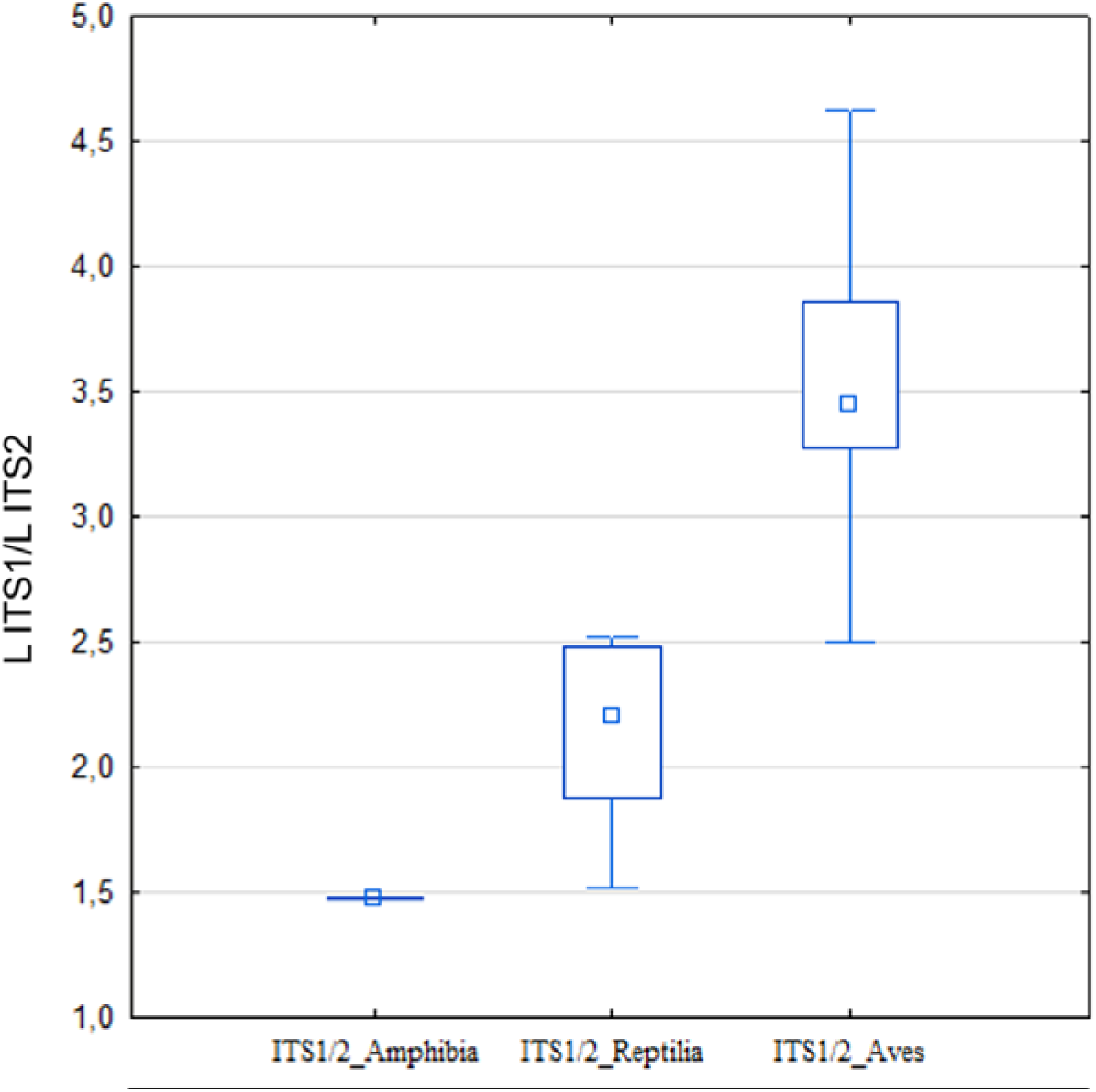
*ITS1* to *ITS2* length ratio in Amphibia (ITS1/2_Amphibia), Reptilia (ITS1/2_Reptilia) and Aves (ITS1/2_Aves).

In Sinapsida evolutionary line the length of both spacers sharply increases in mammals (Fig. 3). In most of the analyzed species *ITS1* is slightly longer than *ITS2* (Fig. 7). In placental mammals, *ITS1* to *ITS2* length ratio is similar to most amniotes, the average being around 1.32. Contrarily to birds and reptiles, in mammals *ITS1* and *ITS2* lengths vary widely even among close orders (Fig. 3, 7).

**Fig. 7.**
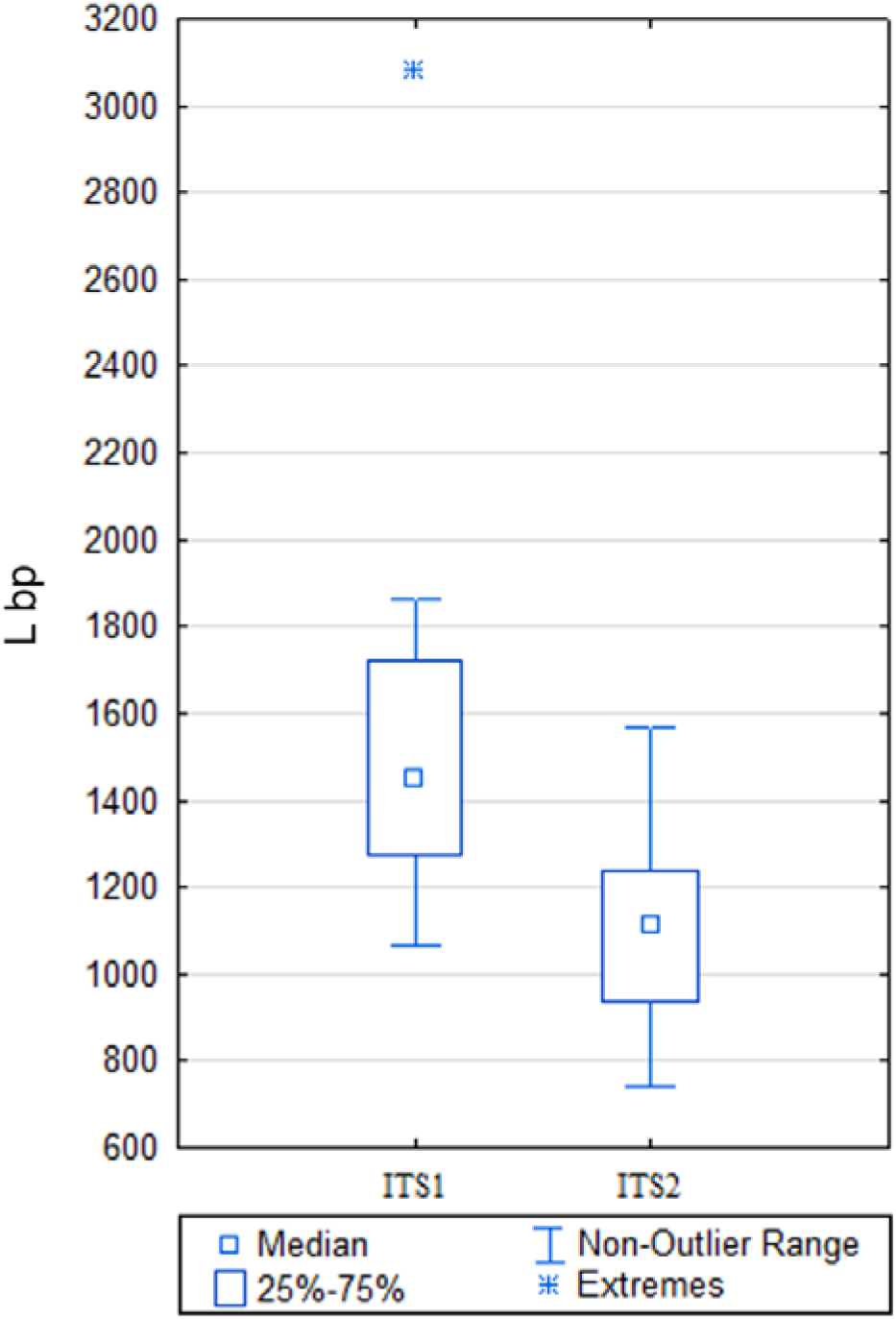
*ITS1* and *ITS2* length range in Mammals.

Marsupial *Monodelphis domestica* constitutes a noteworthy exception. The extraordinarily long *ITS1* (3080 bp) in this species combines with a short *ITS2* of 740 bp. This situation is common to birds and some reptiles, yet not to mammals.

#### 3.2.2. GC pair content variation

GC pair content variation in Sauropsida and Sinapsida evolutionary lines is shown in Figures 8 and 9 respectively. In both charts a smooth GC-content increase can be seen in both *ITS1* and *ITS2* from Ctenophora, Cnidaria, and primitive Deuterostomia to terrestrial vertebrates. In Chaetognatha and Echinodermata, GC pair content is about 61% in *ITS1* and 63% in *ITS2.* In coelacanth (Sarcopterigii), GC pair content in *ITSs* achieves as much as 75 – 80% and increases further to 82–85 *%* in birds and mammals.

**Fig. 8.**
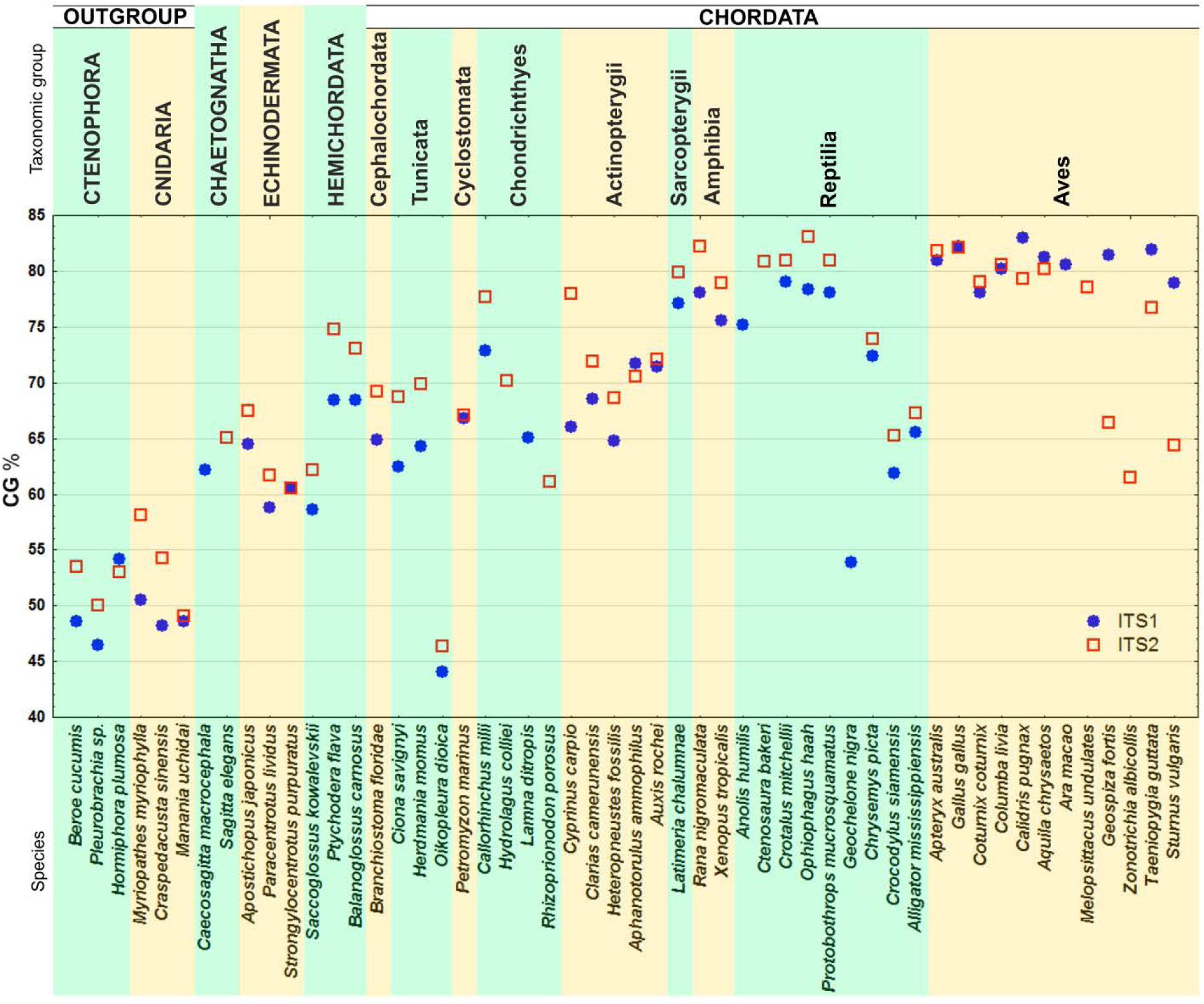
GC-content in *ITS1* and *ITS2* sequences in Sauropsida evolutionary line.

**Fig. 9.**
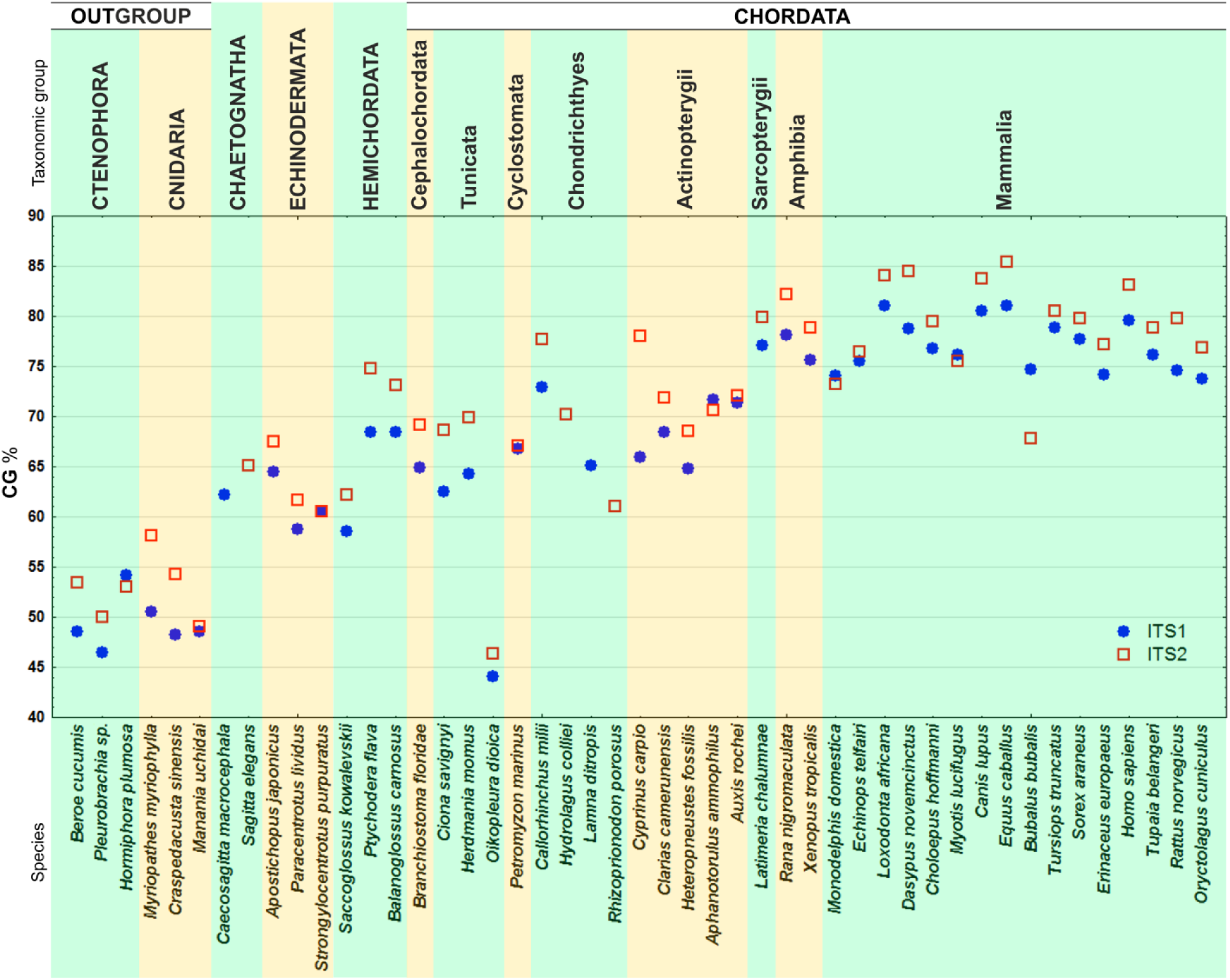
GC-content in *ITS1* and *ITS2* sequences in Synapsida evolutionary line.

Most taxa, with the exception of Aves, feature a higher GC-content in *ITS2* compared to *ITS1* (Fig. 10A). Concurrently, in most taxa GC-content in *ITS1* and *ITS2* is virtually the same within the same species: the difference generally does not exceed 8%, the average being 3.3% and close to 0% in some species (Fig. 10B). Torres et al. (1990) obtained similar results from spacer analysis of some evolutionarily remote eukaryotes. This phenomenon prompts the existence of some concerted evolution patterns stabilizing *ITS1* and *ITS2* nucleotide content in Deuterostomia.

**Fig. 10.**
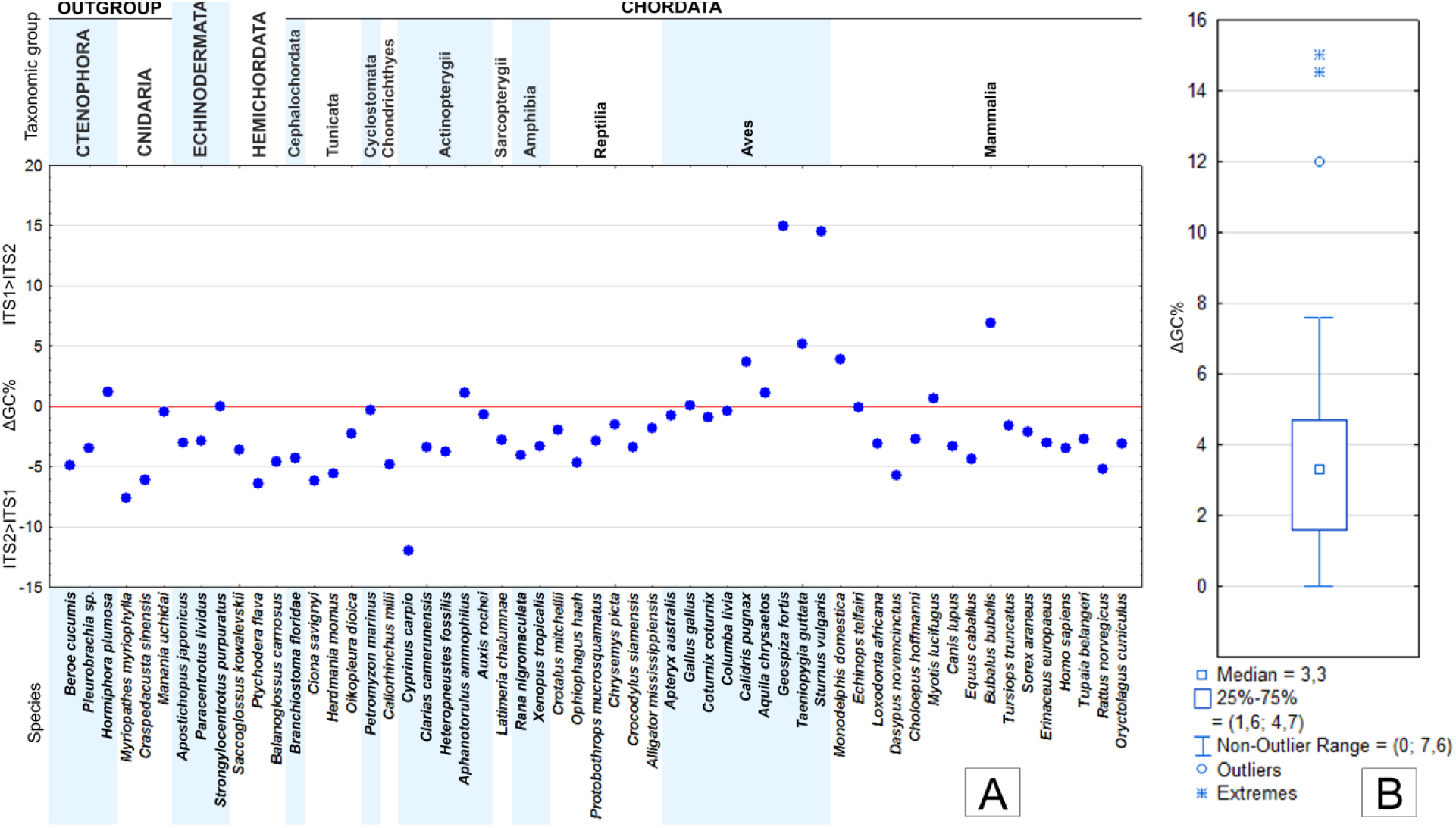
**A.** GC-content variation in *ITS1* and *ITS2* in the analyzed Deuterostomia species. **B.** Box plot of ΔGC% in *ITS1* and *ITS2* in the analyzed Deuterostomia species

#### 3.2.3. Nucleotide compositional shift in the sense DNA sequences of ITS1 and ITS2 in Deuterostomia

The charts in Figures 11 and 12 show nucleotide compositional shift in *ITS* sense of Deuterostomia DNA sequences. The situation observed is generally common to both spacers. Their adenine content is higher than thymine content in the majority of the analyzed species. This feature can already be detected in Ctenophora and Cnidaria members. In Siamese crocodile (*Crocodylus siamensis*) the *ITS1* nucleotide balance shifts towards a higher adenine content. A considerable shift has been recorded only in mammals. In *ITS2* adenine content increase starts with lamprey (*P. marinus*) and is well-marked in most vertebrates. Guanine and cytosine content in *ITS* sense DNA sequences varies insignificantly and virtually stabilizes in painted turtle (*Chrysemys picta*). Therefore, the observed thymine content decrease in *ITS* sense DNA chains correlates to GC-content in double strength DNA of these sequences.

**Fig. 11.**
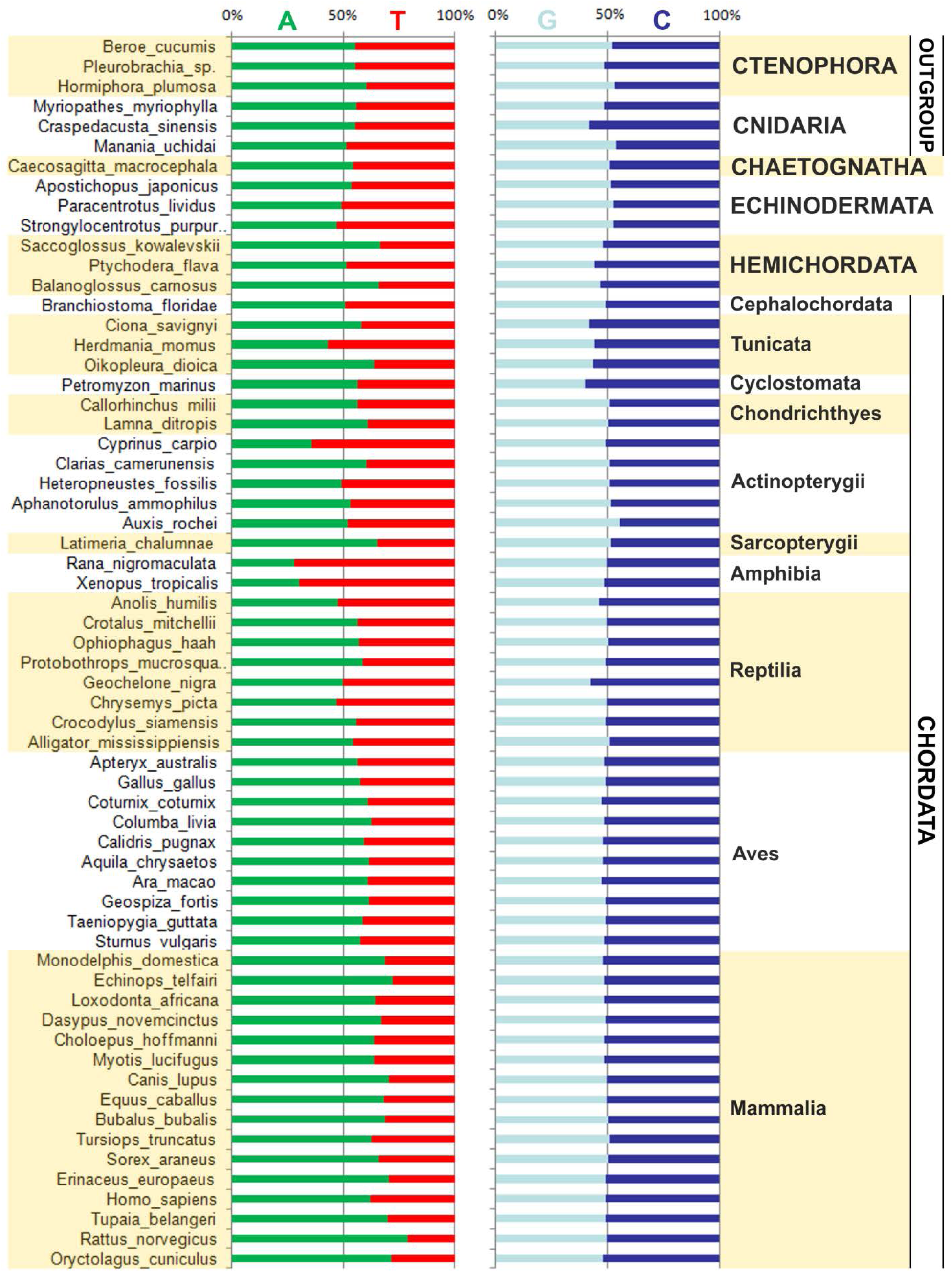
Nucleotide composition shift in *ITS1* sense DNA sequences

**Fig. 12.**
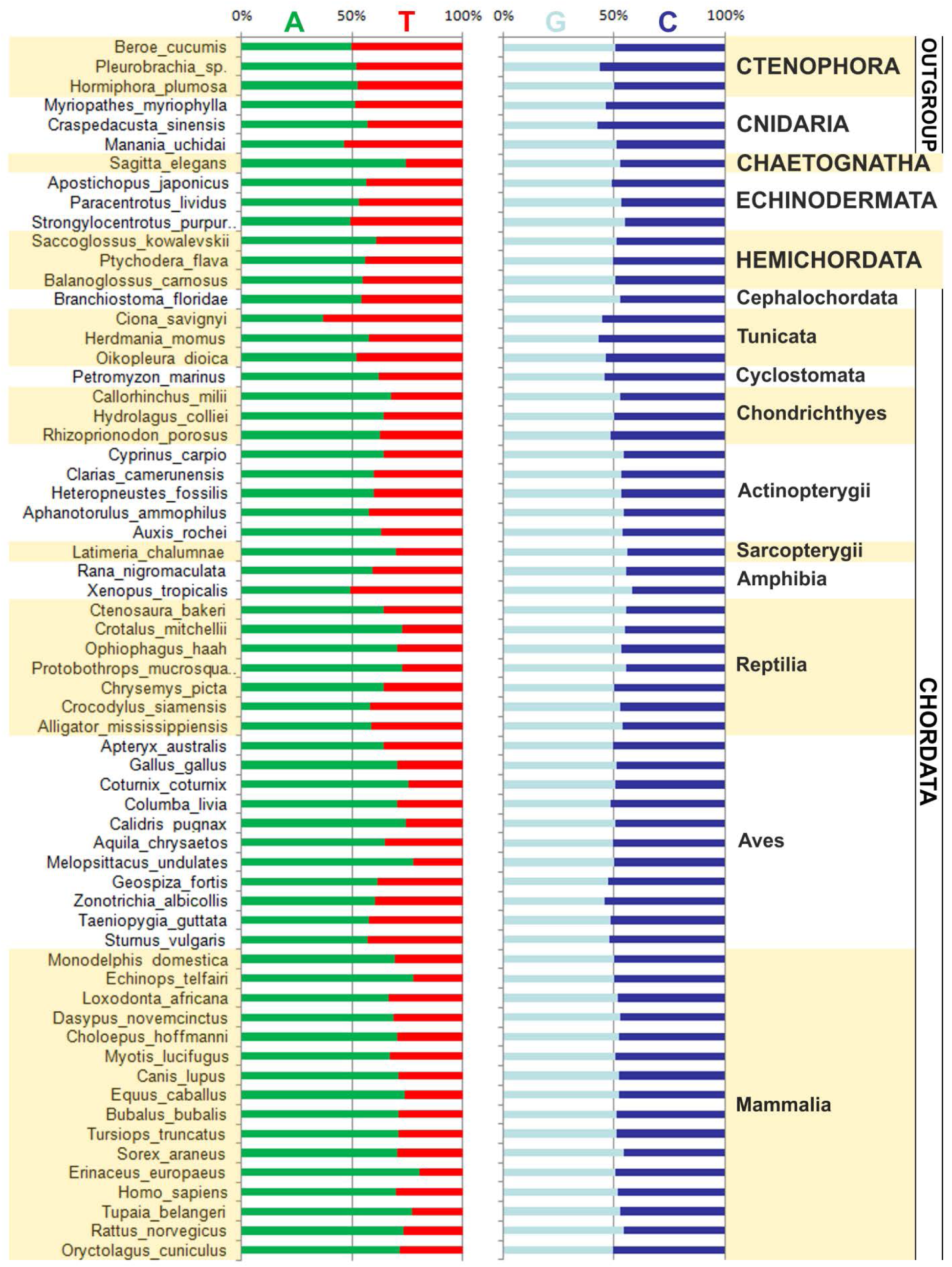
Nucleotide composition shift in *ITS2* sense DNA sequences

Earlier methyl-cytosine deamination in CpG dinucleotides was shown to be involved in plant *ITS1* and *ITS2* alteration towards AT-content increase (Torres et al., 1990). The GC-content increase in Deuterotomia described by these authors is likely to be related to the effect of other mechanisms.

### 3.3. Peculiarities of GC-content in Sauropsida ITS1 and ITS2

The reported common patterns of GC-content variation in Deuterostomia *ITS1* and *ITS2* include a number of exceptions requiring special focus.

In the segment of the chart (Fig. 8) showing GC-content in Sauropsida *ITSs*, we can clearly see a region of GC-content collapse in turtles and crocodiles. GC-content in both *ITSs* is by 20-25% lower in crocodiles (C. *siamensis* and *Alligator mississippiensis)* and 10-15% in turtles (C. *picta)* compared to the rest of Sauropsida. This phenomenon stands out of the general GC increase trend in *ITSs* associated with Deuterostomia evolution. We assume that this phenomenon could be related to expansion of AT-enriched mobile genetic elements (MGC) in spacer sequence. Thus, in Galapagos tortoise (*Geochelone nigra*) *ITS1* featuring a length of 2007 bp and 54 GC-content (NCBI, Nucleotide: AY101758), we have found an AT-enriched fragment of 880 bp (Supplementary File 2). The similarity of this fragment to non-LTR retrotransposon CR1-4 CPB sequence achieves 93%. NCBI database contains 15 complete *ITS1* sequences of five *Geochelone* species (NCBI, Nucleotide: AY101749–AY101763) including AT-enriched fragments that are also homologous to CR1-4 CPB. The repetitiveness of such cases excludes the probability of an artifact. MGE presence in *ITSs* Deuterostomia sequences has not been reported earlier. MGE insertion is likely to be one of *ITSs* evolutionary patterns. MGE insertions have not been found in the *ITSs* of other analyzed Deuterostomia taxa. The homology of insertions and source MGE sequences could have been terminated due to the high evolutionary rate of rDNA spacers (Wang et al., 2015).

In birds, from ruff (*Calidris pugnax*) and onwards, GC-content in *ITS2* decreases compared to *ITS1* (Fig. 7), which is uncommon to the great majority of the analyzed taxa. In Passeriformes, the most numerous Aves order, GC-content in *ITS2* is 1.5 times lower than that of other orders. For example, GC-content in zonotrichia (Zonotrichia *albicollis*) *ITS2* is 61,5%, while in other orders its can achieve as much as 80%. GC-content in *ITS1* exceeds *ITS2* by 15% in medium ground finch (*Geospiza fortis*) (Fig. 9). *ITS2* length in Passeriformes increases compared to the rest of the analyzed Aves, subclass Neognathae by at least 132 bp (Fig. 2). The insertions accounting for this increase are unique to Passeriformes and AT-enriched in 3 out of the 4 analyzed species. It is quite possible that MGE insertion could have caused variations in GC-content in Passeriformes *ITS2* and in Galapagos tortoise.

## 4. Discussion

Our findings allow suggesting that internal transcribed spacer structural variation from inferior to superior taxa of Deuterostomia is associated with four key aspects: a) spacer length increase; b) GC-content increase in both *ITSs;* c) uniform GC-content in *ITS1* and *ITS2* within the same species; d) thymine content decrease in sense DNA sequences in both *ITSs*.

The most considerable *ITS*s extension has been identified in superior vertebrates and members of subclass *Elasmobranchii* (5–12 fold versus the members of the rest of the analyzed taxa). Superior vertebrates and *Elasmobranchii* are not sister taxa and have been evolving independently of each other. Convergent extension of *ITSs* of their members could be related to various selection factors. Both of these groups strongly differ ecologically and physiologically, so that physical environmental factors and the rate of metabolic processes appeared to have been of no key role in such changes.

*ITSs* length extension must have been related to the impact of focused selection, yet the true causes and patterns of this extension remain unclear. It would be feasible to analyze spacer length extension in the context of nucleotide composition shift. Spacer differences between close species of Deuterostomia are primarily defined by indels of several nucleotides or large fragments (of 20 or bp or longer). The possible role of MGE insertion in *ITSs* extension cannot be excluded either.

Contrarily to more primitive taxa of Deuterostomia, spacer extension within Amniota group evolved in two directions: in placental mammals both spacers generally extended commensurately, while in marsupials, reptiles and birds spacer length ratio shifted towards significant extension of *ITS1* (2–3 fold). Probably, a similar disproportion could have been discovered in the common amniotic ancestor of Sauropsida and Sinapsida, which is specifically evidenced by the obviously disproportional spacer lengths of marsupial mammals.

In the evolutionary line of Deuterostomia *ITSs* length extension is accompanied by *ITSs* nucleotide composition shift. Specifically, GC-content smoothly increases: the difference between primitive Deuterostomia (Chaetognatha and Echinodermata) and *Tetrapoda* and *Sarcopterygii* is about 15 – 20%. Concurrently adenine content increases, while thymine content respectively decreases in *ITS* sense DNA sequences. The most likely pattern underlying this nucleotide composition shift could consist in GC-biased gene conversion (gBGC) leading to GC-content increase for the account of gene conversion in the course of recombination and selective repair of the appearing non complementary TG pairs to CG (Fryxell and Zuckerkandl, 2000; Galtier et al., 2001; Duret and Galtier, 2009). gBGC phenomenon has been noted in a wide range of eukaryotes (Pessia et al., 2012), ectopic genetic conversion presumably being one of the core patterns of concerted evolution of rDNA repeat units (Naidoo et al., 2013). We could assume that the observed GC-content increase and thymine content decrease in Deuterostomia *ITS* with evolution is associated with the increasing gBGC capacity and/or impact of selection focused on GC pair binding in spacer sequences. The same processes could also account for sustaining GC/AT content similarity between internal transcribed spacers on intra-species basis.

*ITS*s length extension and the related GC-content increase lead to formation of infusible secondary structures containing multiple hairpins in pre-rRNA transcripts (Dyomin et al., 2016). The evolutionary effect of this spacer transformation remains unclear. Presumably, secondary structures are better recognized by pre-RNA splicing factors, which contributes to ribosome RNA formation. Furthermore, spliced pre-rRNA fragments enriched with G- and C-bases produced an enormous amount of double-stranded RNAs (dsRNA), whose further destiny is still unclear. Pre-rRNA synthesis rate has been reported to be consistently high in interphase nucleus and change considerably in the course of cell division (McStay and Grummt, 2008; Brown and Szyf, 2008). It is quite possible that in vertebrates, taking into account their advanced genetic chains, dsRNA forming from *ITS* transcripts could be one of the components of cell cycle genetic regulatory pattern.

## Abbreviations

HTS: high-throughput sequencing
IGS: intergenic spacer
ITS: internal transcribed spacer
WGS: whole genome sequence shotgun
SRA: sequence read archive
MGC: mobile genetic elements
gBGC: GC-biased gene conversion

## Acknowledgements

The research has been financed by Saint-Petersburg State University (project number 1.50.1043.2014) and Russian Basic Research Foundation (project 15-04-05684). The work was performed on the base of «Chromas» resources centre of Saint-Petersburg State University Scientific Park.

### Appendix A. Supplementary material

Supplementary File 1

All *ITS1* and *ITS2* sequences used in this study

Supplementary File 2

Insertion of non-LTR retrotransposon CR1-4 CPB into *Geochelone nigra ITS1* sequence

